# Rhythmically bursting songbird vocomotor neurons are organized into multiple sequences, suggesting a network/intrinsic properties model encoding song and error, not time

**DOI:** 10.1101/2023.01.23.525213

**Authors:** Graham C. Fetterman, Daniel Margoliash

## Abstract

In zebra finch, basal ganglia projecting “HVC_X_” neurons emit one or more spike bursts during each song motif (canonical sequence of syllables), which are thought to be driven in part by a process of spike rebound excitation. Zebra finch songs are highly stereotyped and recent results indicate that the intrinsic properties of HVC_X_ neurons are similar within each bird, vary among birds depending on similarity of the songs, and vary with song errors. We tested the hypothesis that the timing of spike bursts during singing also evince individual-specific distributions. Examining previously published data, we demonstrated that the intervals between bursts of multibursting HVC_X_ are similar for neurons within each bird, in many cases highly clustered at distinct peaks, with the patterns varying among birds. The fixed delay between bursts and different times when neurons are first recruited in the song yields precisely timed multiple sequences of bursts throughout the song, not the previously envisioned single sequence of bursts treated as events having statistically independent timing. A given moment in time engages multiple sequences and both single bursting and multibursting HVC_X_ simultaneously. This suggests a model where a population of HVC_X_ sharing common intrinsic properties driving spike rebound excitation influence the timing of a given HVC_X_ burst through lateral inhibitory interactions. Perturbations in burst timing, representing error, could propagate in time. Our results extend the concept of central pattern generators to complex vertebrate vocal learning and suggest that network activity (timing of inhibition) and HVC_X_ intrinsic properties become coordinated during developmental birdsong learning.

## Introduction

Network activity driving experience-dependent synaptic plasticity and activity-dependent regulation of cellular intrinsic properties (IPs) are two mechanisms contributing to learning and memory phenomena (Feldman, 2009; Mozzachiodi and Byrne, 2010; Daou and Margoliash, 2021). Recent studies suggest how both contribute to memory stabilization (Titley et al., 2017; Grasselli et al., 2020; Titley et al., 2020). One potential mechanism for integrating network activity with plasticity of IPs is via spike rebound excitation, where the hyperpolarization activated inward current and the T-type Ca^++^ current regulate the timing, strength and duration of the rebound burst upon release from inhibition, resulting in rapid depolarization and rebound spiking. The timing of modulation of inhibition depends on network activity. While there is substantial evidence gathered in vitro in support of this model, directly correlating this mechanism with behavioral plasticity is challenging (Alviña et al., 2008; Person and Raman, 2012; Hasselmo, 2014; Reato et al., 2016).

Bird song is an advantageous vertebrate system for examining the interactions of network and cellular substrates of learned motor production. Within the telencephalic nucleus HVC, the HVC_RA_ neurons project to RA (primary motor cortex analog), the HVC_X_ neurons project to Area X (basal ganglia), and there may be a small number of projection neurons (PNs) that project to both targets (Kornfeld et al., 2017; Benezra et al., 2018). The adult songs of zebra finches are highly stereotyped and thus present complex but highly spectrotemporally structured behaviors. During singing, HVC PNs emit phasic bursts of spikes (≈ 10 ms duration) whereas HVC interneurons (HVC_INT_) are tonically active with local minima and maxima in their firing patterns. Of those PNs that spike during the song motif that comprises the canonical sequence of syllables, HVC_RA_ emit one spike burst and HVC_X_ emit one or more spike bursts. Each HVC_RA_ and HVC_X_ spike burst is precisely timed to a specific moment in the motif. Whether the times when PNs burst is related to features of song remains unresolved (Hahnloser et al., 2002; Amador et al., 2013). In one conceptualization of the functional organization of HVC, the burst times from all the HVC_RA_ and HVC_X_ contribute to a single sequence distributed continuously across the motif, explicitly representing time itself, with little (Lynch et al., 2016) or no (Picardo et al., 2016) representation of song features. This is the so-called “clock” model of HVC activity.

HVC_X_ neurons participate in auditory processing mechanisms central to juvenile birdsong learning (Roberts et al., 2010), and exhibit changes in dendritic spines and increases in intrinsic excitability after deafening (Tschida and Mooney, 2012). Recent results demonstrate that the IPs of HVC_X_ within each bird are similar to each other, with the variation observed from bird to bird related to how similar are the songs of the birds (Daou and Margoliash, 2020). Thus, descriptive features of song correlate with the distribution and variance of features of HVC_X_ IPs. We wondered how this would be expressed at the network level, intermediate between cellular properties and behavior. The principal feature describing the activity of HVC PNs during singing is the timing of when the spike bursts are released (Hahnloser et al., 2002; Kozhevnikov and Fee, 2007). We reasoned that the bird-specific distributions of HVC_X_ IPs should bias the timing of HVC_X_ bursting during singing towards bird-specific patterns, if those bursts arise from the action of spike rebound excitation (Daou et al., 2013).

## Results

Lynch et al. (2016) presented a large data set of high impedance extracellular recordings collected during singing from five adult zebra finch. After extensive preliminary analysis of the veridical burst time data from Bird 1 (“Bird 2” of Kozhevnikov and Fee (2007), available to us from data previously provided to DM by A. Kozhevnikov) and approximations of the data from Birds 2-5 that we extracted from the Lynch et al. (2016) pdf, we then re-analyzed the Lynch et al. (2016) data, provided by the authors. For a brief summary of the relevant data collection procedures refer to Materials and Methods, and for more detail refer to Lynch et al. (2016).

The Lynch et al. (2016) data includes “multiburst” neurons emitting two or more bursts per motif (Supplemental Table 1). HVC multiburst neurons are reported to be HVC_X_ as identified using antidromic stimulation techniques (Kozhevnikov and Fee, 2007; Lynch et al., 2016). In a population of 105 identified HVC_X_ recorded from 7 adult birds, 18 emitted zero bursts per motif, 45 emitted one burst per motif, and 42 emitted between 2–4 bursts per motif (Kozhevnikov and Fee, 2007).

The Lynch data set was strongly biased towards HVC_X_ (Supplemental Table 1). For PNs that were confirmed (that is, identified by antidromic stimulation) HVC_X_ were 62.5% of Bird 4 neurons and ≥ 80% for the other birds. Across the five birds, of all positively identified PNs, positively identified HVC_X_ contributed 66–93% of all bursts. In what follows we mostly analyzed just the HVC_X_ data, with a focus on multiburst neurons. Expanding the data set to include putative PNs (see Methods), multiburst HVC_X_ were 42-65% of all HVC_X_ neurons in the five birds. Bursts from multiburst neurons dominated the data set for the three birds with the larger samples (67%, 73%, 63%, Birds 1-3, respectively) and also contributed substantially for the other two birds (36%, Bird 4; 39% Bird 5). Since some of the multiburst neurons includes putative PNs (neurons that had characteristic features of HVC PNs but were not identified by antidromic stimulation; see Lynch et al. (2016)), the sample we analyzed was highly enriched in HVC_X_ but may include very small numbers of other as yet poorly defined cell classes (see Methods). In the rest of the Results, for ease of exposition, we refer to these data as HVC_X_.

### HVC_X_ burst rhythmically during singing

We observed that multiburst HVC_X_ evinced simple low-order structure in the timing of the bursts. To explore this, using the Lynch et al. (2016) definition of a spike burst, first we examined the timing of intervals between bursts. To this end we constructed histograms (10 ms bins) from the pooled inter-burst intervals (IBIs) from each multiburst neuron within a bird. Birds 1 and 5 had relatively flat histograms; the Bird 3 histogram had at least two peaks, and the histograms of Birds 2 and 4 were dominated by a single peak. The position of the peaks varied from bird to bird. Simple scaling or truncation of the time axes (to account for different motif durations) could not account for these differences. Thus, there was evident first-order structure in the timing of HVC_X_ multiple bursts and variation from bird to bird. This also demonstrates that within each bird the burst intervals of multiburst HVC_X_ neurons were very highly correlated (statistically dependent).

A subset of PN spikes did not qualify as contributing to bursts, and the IBI histograms capture a first order statistic (and with a somewhat arbitrary bin width). To address these limitations, we also evaluated the spike firing rate of each multiburst PN, thus including all the spikes of each neuron in the analysis. Autocorrelograms of the firing rates were created from the 1 ms-binned spike rates for each neuron. An example is shown in Fig. 2A. The mean spike rate of a neuron (Figure 2A, top panel) is autocorrelated yielding a principal peak at zero lag and burst-interval-related peaks at positive and negative lags (Figure 2A, middle panel). Focusing just on the positive lags we defined the numerically largest values as the peaks (Figure 2A, bottom panel).

To search for common second order patterns across the neurons of each bird, we then scaled the autocorrelogram height of each neuron on the unit interval and summed them, resulting in a single plot per bird (Figure 2, Birds 1–5). These summed autocorrelograms show distinct peaks within a given bird, indicating that the intervals between periods of high firing rate were similar across neurons within a bird. Features of the summed autocorrelograms can be related to features of the corresponding PSTHs (Figure 1), although the autocorrelograms have more detail.

**Figure 1.**
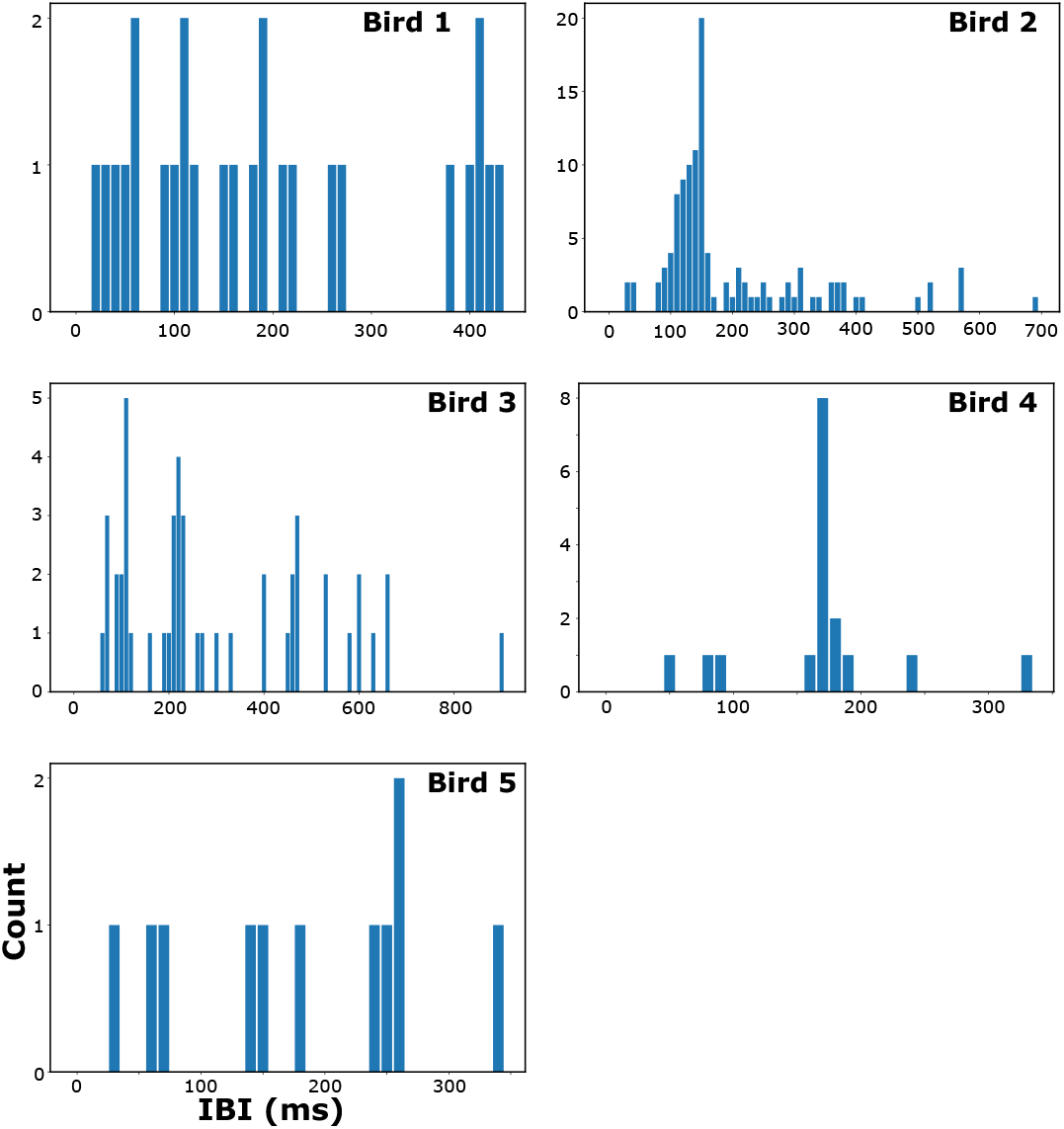
Inter-burst interval histograms of multiburst HVC_X_. The distributions vary among the birds, with no peaks evident for Birds 1 and 5, clear single peaks evident for Birds 2 and 4, and several peaks evident for Bird 3. 10 ms binwidth, all histograms.

**Figure 2.**
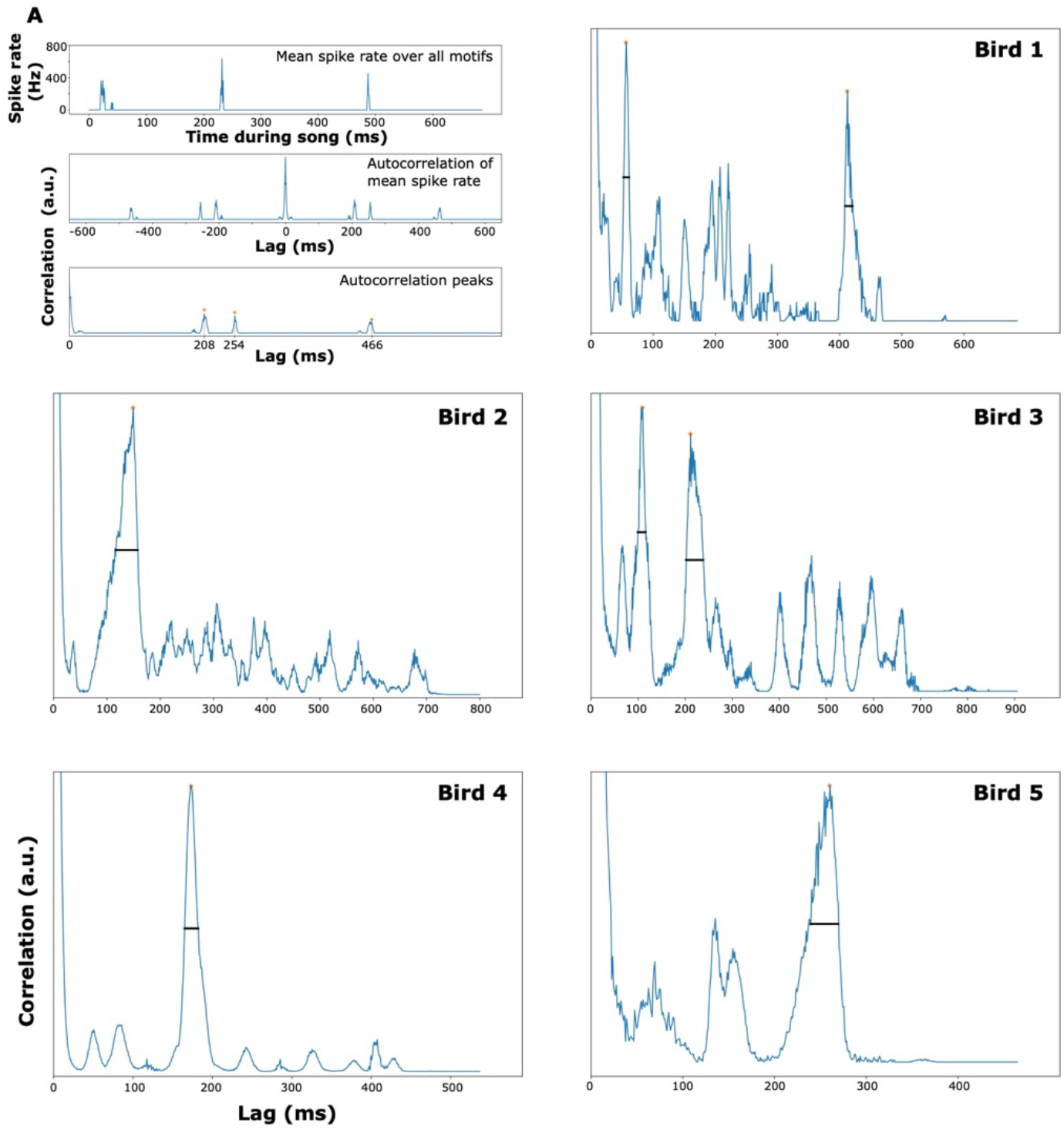
Autocorrelations of multiburst HVC PNs. (A) Example autocorrelation analysis for neuron 25 of Bird 1. Top panel: 1 ms-binned spike rate averaged over all motifs. Middle panel: autocorrelation of averaged spike rate for lags up to the motif duration. Bottom panel: positive domain of autocorrelation, with peaks indicated by orange stars. (Birds 1–5) Summed autocorrelograms for each bird. Per-neuron autocorrelograms as in (A), bottom panel, were normalized to the unit interval and summed to produce these plots. To give context, the number of neurons contributing to the top 50% (black horizontal line) of each peak relative to the total number of multiburst neurons is Bird 1, 28% (5/18) and 22% (4/18) (first and second peaks); Bird 2, 74% (50/68); Bird 3, 23% (10/44) and 39% (17/44) (first and second peaks); Bird 4, 93% (13/14); Bird 5, 60% (6/10).

The structure of the summed autocorrelograms show clear differences between birds. To test the significance of these differences we computed the zero-lag correlations (dot products) between the autocorrelograms of all pairs of neurons within each bird and between all pairs of neurons across each pair of birds. Some of the resulting distributions were not normally distributed, so we assessed the differences in the distributions using the K-S statistic. In 18/20 comparisons the distributions were statistically significantly different, with only Bird 5 self vs. cross of Bird 4 and Bird 5 self vs cross of Bird 3 failing to achieve significance after Bonferroni correction (Supplemental Table 2). (Bird 5 had the smallest data set.) Thus, the functional properties of multiburst neurons varied among the birds. We note that individual differences in song is a hallmark of birdsong learning.

### HVC_X_ bursts during singing are organized in bird-specific multiple sequences

The similarity of burst timing in the HVC_X_ within each bird for neurons that are recruited at different moments of song should constrain the dynamics of the population bursting. Specifically, it implies that later bursts should be organized into sequences at time offsets relative to the initial bursts. Lynch et al. (2016) (their Figure S1) provided a useful representation for examining the structure of the population of HVC activity in relation to the song motif. In that “neuron cascade” representation, bursts from the same neuron are plotted on the same row, and neurons are sorted from top to bottom by the time of their first burst relative to the canonical song motif. Preserving the ordinal relationship of the timing of the first bursts while retaining the neuron identity of the subsequent bursts aligns all bursts in relation to when they occurred during the song motif while representing the flow of neurons as they were initially recruited during the motif.

Lynch et al. (2016) evaluated HVC_X_ and HVC_RA_ together. Here, examining such representations but only for the HVC_X_ for each of the five birds (Figure 3A–E; first bursts in blue, later bursts in orange), we observed a clearly defined approximately linear slope of initial bursts (or piecewise fits where appropriate), and quasi-linear segments of later bursts parallel to the slope of the initial burst. A striking example is found in Bird 4, with over 50% (10/17) of later bursts arranged around a single line (Figure 3D, dotted line). Other clear example are the later bursts of Birds 1 and 5 (Figures 3A, 3E, dotted lines). We stress that here we have placed the dotted lines manually, to draw attention to some of the more obvious examples of this structure. To quantify the apparent structure of later bursts, we developed a procedure to analyze the significance of clustering of subsequent bursts (Supplemental Figure 1). First, we projected the later bursts onto an axis normal to the first burst line (Supplemental Figure 1A, B). Then, we tested whether the bursts cluster on this axis, and confirmed that they do using the Ripley (1976) L statistic (Supplemental Figure 1C–H).

**Figure 3.**
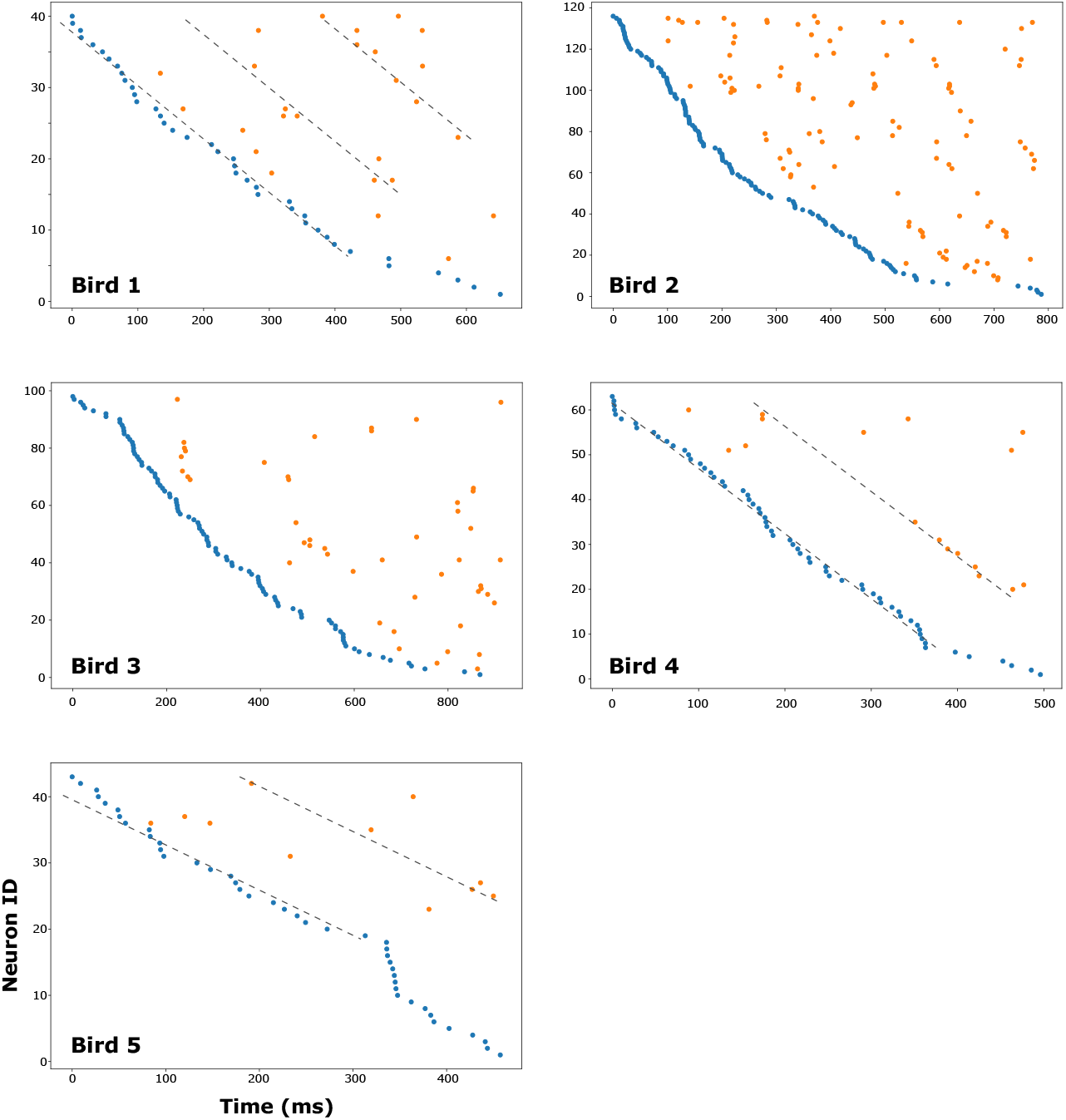
Neuron cascade representations reveal structure in the timing of later bursts. Burst times of HVC_X_ PNs for each bird, with all the bursts of each neuron on one row and neurons ordered vertically by the time of their first bursts within the song. First burst times are in blue, later burst times in orange. For Birds 1, 4, and 5 dotted lines with slopes fit to the corresponding initial slope lines show examples of bursts that cluster at a fixed temporal offset to the initial bursts.

While these results demonstrate that there are multiple sequences of HVC_X_ bursts that unfold over the time course of a song motif, there is no *a priori* reason to believe that a burst cascade should be well-characterized by a linear fit. Therefore, we developed an analysis to examine the arrangement of subsequent bursts without the parameterization imposed by the linear regression and normal-axis projection method. Starting with a neuron cascade representation for a given bird, e.g., Bird 1 (Figure 4A), we shifted each neuron’s burst times so that its first burst occurred at zero, while preserving all within-neuron IBIs (Figure 4B, top panel). All later bursts could then be projected onto the time axis (Figure 4B, bottom panel), restricting subsequent analysis to this one-dimensional representation. Bursts close together on this axis will be arranged in the original burst time plots in such a way as to follow a similar progression in time as the first bursts. Analyzing these data with the Ripley’s L metric we found that Birds 1-4 displayed clustering at short timescales that lay outside the 95% confidence band for the surrogate data. Bird 5 did not exhibit clustering greater than that displayed by its surrogate data, but this is likely explained by its small set of later bursts (n = 11).

**Figure 4.**
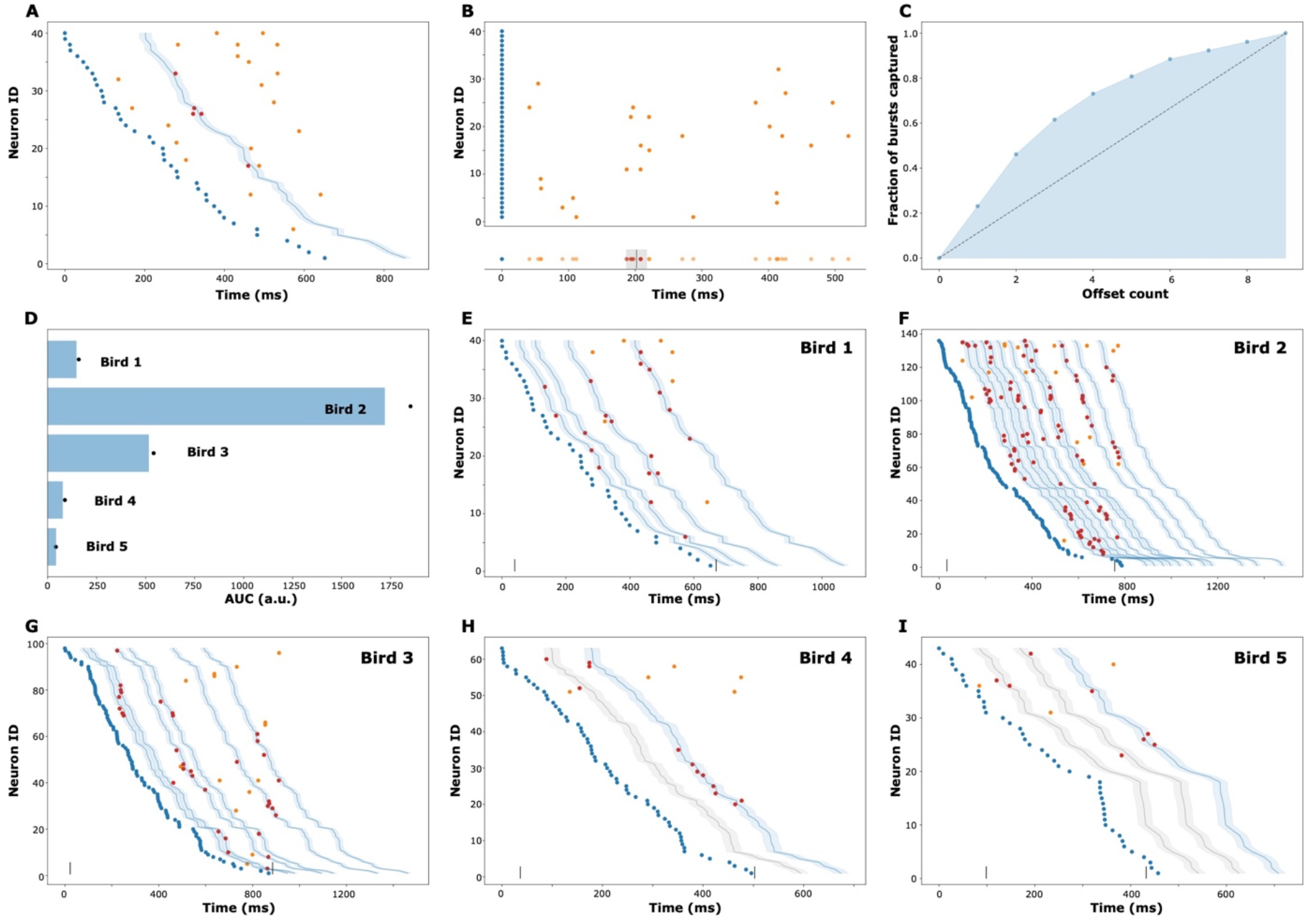
Multiple Sequences of Later Bursts During Singing. For all panels, initial burst times are in blue, while later burst times within and outside of capture windows are in red and orange, respectively. All results are from the larger capture set algorithm (see text). (A) Burst times for Bird 1, plotted as in Figure 3A. An example offset is at 202 ms (dark blue line at center of shaded 30 ms capture window). (B) Top panel: burst times for Bird 1, adjusted per-neuron so that each neuron’s first burst occurs at 0, preserving within-neuron IBIs. Bottom panel: projection of all bursts from the top panel onto the time axis. An offset at 202 ms (black vertical line) captures spikes in a ±15 ms surrounding region (grey shading). (C) The cumulative proportion of later bursts captured by increasing numbers of offset regions for Bird 1. Proportion of later bursts captured is depicted by blue points, with the shaded blue region the area under this curve (AUC). The diagonal black line shows the capture result if every offset had only captured a single burst. (D) Summary of comparable AUC analysis for each bird. The blue bars denote the 95% confidence interval for the surrogate random uniformly-distributed data. Black points represent the AUC for the real data. (E–I) All capture regions for Birds 1-5. The blue offset regions are those which capture three or more later bursts. For Birds 4 and 5, additional offsets which captured two bursts are plotted in gray. The two vertical bars on each time axis denote the beginning and end of each motif.

Having demonstrated the existence of clustering of later bursts we then determined how many clusters existed and the timing of those clusters for each bird. The question can be seen to be equivalent to a well-known problem in combinatorics, the set cover problem, with a set defined by an interval on the time axis containing one or more burst times. The one-dimensional geometric version of this problem admits of a simple greedy solution (see Methods). We applied this algorithm to the burst locations on the time axis to generate a collection of offsets, together with the later bursts that they capture. A given offset captured all later bursts that fell within a fixed window of the offset. When mapped back onto the original burst time plots, this set of bursts can be seen to fall along a portion of an offset first burst line (Figure 4A). The examples shown in Figure 4 all used a window of 30 ms centered at each offset. We confirmed that they are representative of the effects seen by using a range of window sizes from 20 ms to 60 ms. Each bird’s later bursts were covered by a collection of offsets smaller than the number of bursts, often dramatically so (Bird 1: 9 offsets captured all 26 later bursts; Bird 2: 20 offsets captured all 113 later bursts; Bird 3: 16 offsets captured all 48 later bursts; Bird 4: 7 offsets captured all 17 later bursts; Bird 5: 6 offsets captured all 11 bursts).

To evaluate the statistical significance of these results we compared the performance of the set cover algorithm on real data against its performance on the surrogate uniformly-placed data. We constructed a plot for a set cover solution, similar to a scree plot (but inverted around the unit slope line), of the proportion of the later bursts captured versus the number of offsets, once they had been sorted in descending order by proportion of later bursts captured. A given solution will describe a monotonically increasing curve, whose slope is everywhere nonincreasing (Figure 4C). To produce a single value for comparison between the real and surrogate data, we took the area under this curve (AUC). For Birds 1-4, the AUC for the real data was significantly greater than that for the surrogate data (Bird 1, p=0.042; Bird 2, p=0.0002; Bird 3, p=0.0023; Bird 4, p=0.0001; p-values derived via bootstrap). The set cover solution for Bird 5, which had the fewest later bursts, did not rise to significance (p=0.11). With Bonferroni correction for five birds, Bird 1 becomes non-significant. These values are largely insensitive to the size of the window.

The set cover solutions described above capture all of a bird’s later bursts using the fewest offsets but ignores biologically significant solutions in which some offsets capture more bursts. We therefore extended the analysis above using a modified algorithm that preferentially selects large capture sets (see Methods), rather than minimizing the total number of sets. For this algorithm a smaller number of offsets covered most of each bird’s later bursts than for the prior algorithm (Bird 1: 4 offsets captured 19/26 later bursts; Bird 2: 11 offsets captured 96/113 later bursts; Bird 3: 6 offsets captured 34/48 later bursts; Bird 4: 2 offsets captured 12/17 later bursts; Bird 5: 3 offsets captured 8/11 bursts). The same pattern of values obtained, with Birds 1-4 having significantly greater AUC than their surrogates, and Bird 5 not being significantly different from its surrogates (Bird 1, p=0.0032; Bird 2, p=0.0001; Bird 3, p=0.0016; Bird 4, p=0.0001; Bird 5, p=0.14; p-values derived via bootstrap) (Figure 6D). Birds 1–4 retain significance following Bonferroni correction for five birds. These values, too, were insensitive to the choice of half-window size.

**Figure 5.**
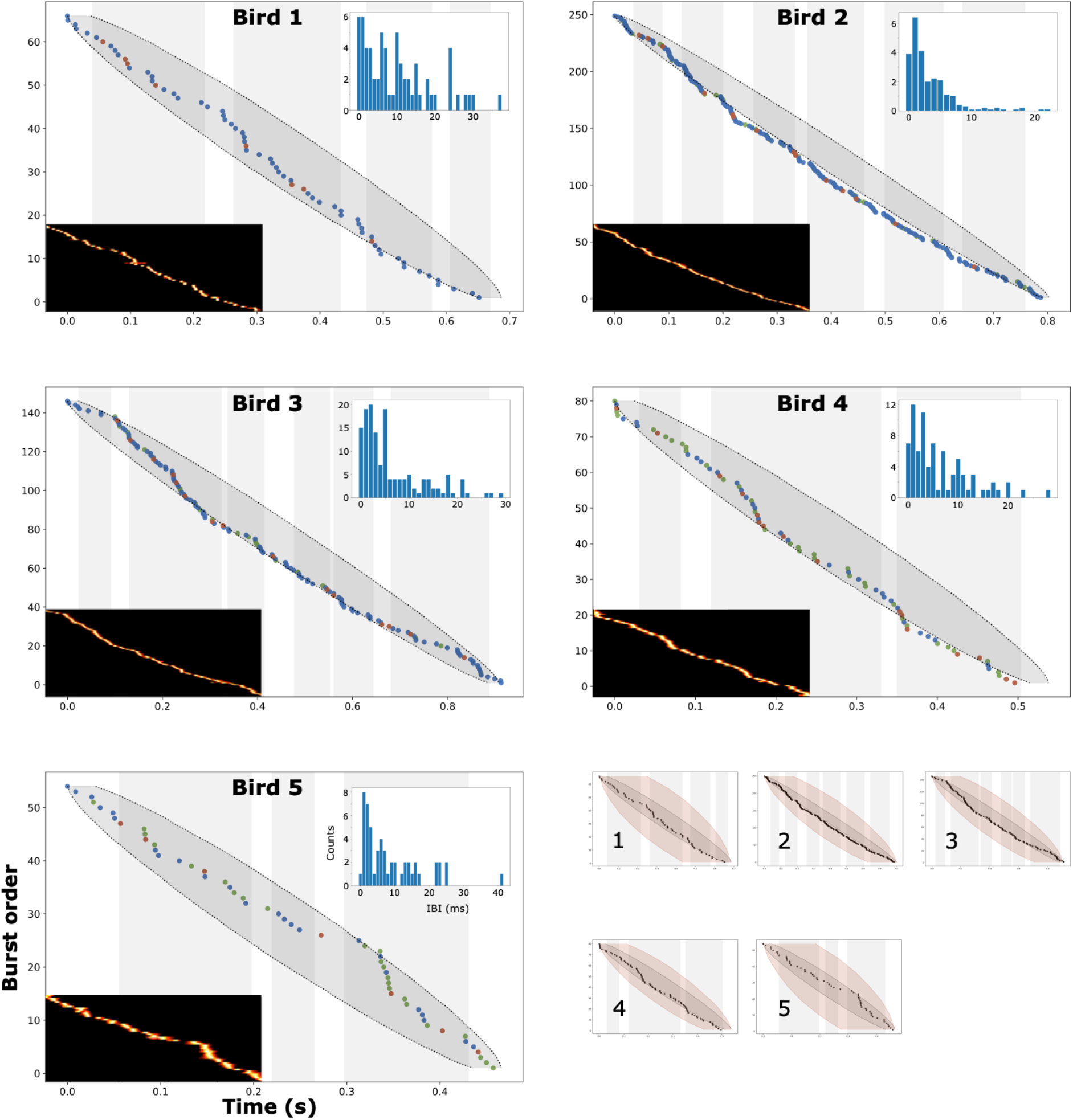
Burst cascade representations of population HVC_X_ activity during singing. (Birds 1–5) Burst times for each bird (HVC_RA_ in red, HVC_X_ in blue, putative PNs in green), with bursts pooled across neurons and ordered vertically by time within the song.. Syllables are marked by shaded gray columns. The dark gray shaded region denotes where 95% of random uniformly distributed surrogate datasets fall. Insets in lower left are of the same bursts as plotted in Figure 1 of Lynch et al. (2016). Insets in top right are IBI histograms computed for the same data. (Bottom right) Each bird is identified by number. All bursts from all PNs are represented by black dots with confidence intervals in grey. The red shaded region denotes the 95% confidence band for random uniformly distributed surrogate datasets of the same size as the positively-identified HVC_RA_ dataset for each of the birds.

**Figure 6.**
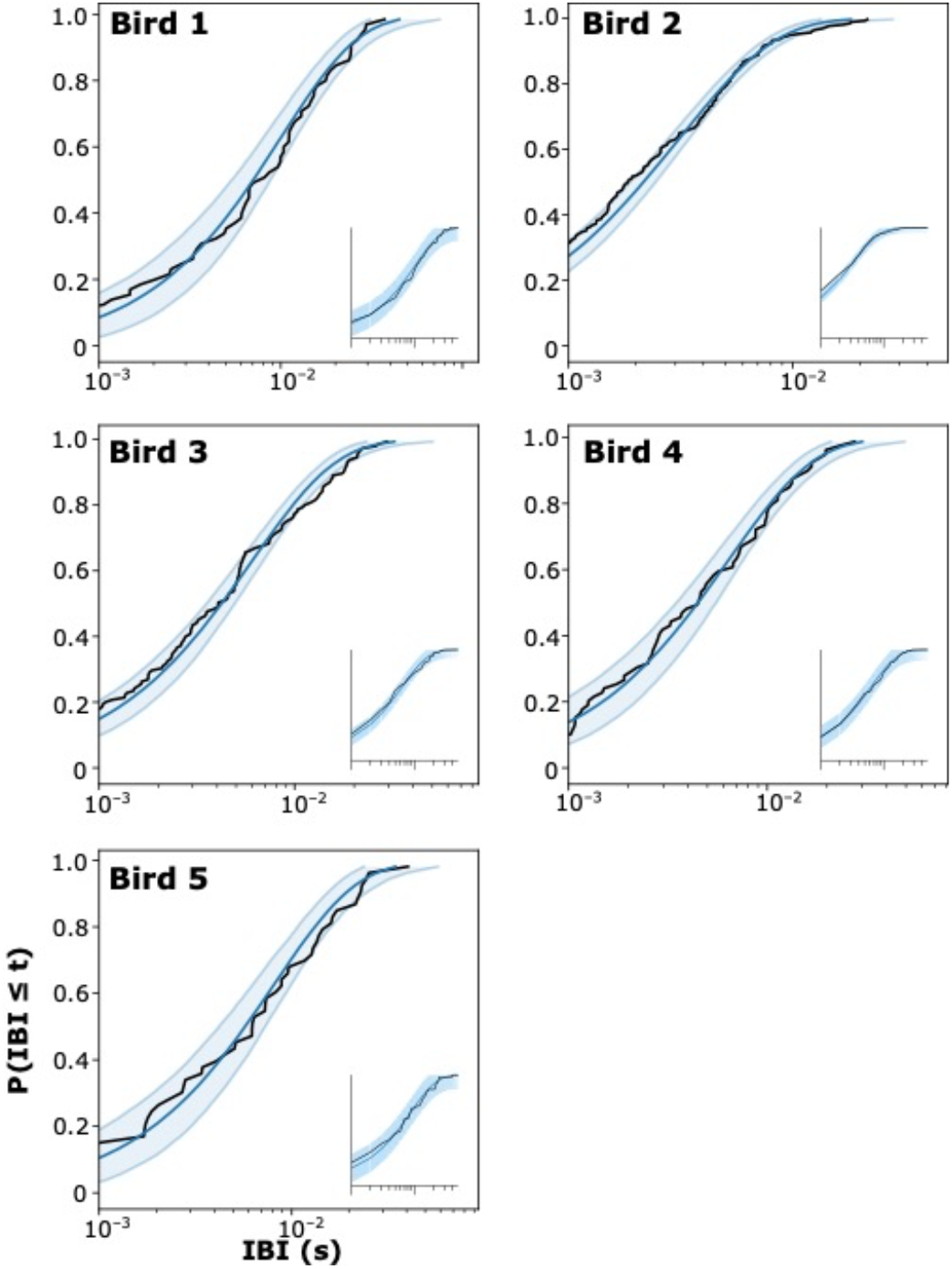
Cumulative density functions of IBIs of HVC projection neurons. Cumulative density functions for the HVC projection neuron IBI distribution for each bird. Real data denoted by black lines, surrogate data by dark blue line (median for surrogates) and light blue shaded region (95% confidence interval). The original CDFs from Lynch et al (2016) for each bird are displayed as insets.

The offsets capturing three or more bursts for all birds are plotted in Figures 4E–4I, providing visual confirmation of the fits described above. It is immediately apparent that different offsets are emphasized in different birds. For these imputed sequences at any given moment in song there can be multiple sequences engaged simultaneously – bursts occurring later in one sequence occurring at the same time interval as bursts occurring earlier in another sequence. Such overlap is also observed directly in the data for all three birds with the larger numbers of multiburst neurons (Birds 1–3). For example, for Bird 3 different bursts occurring at roughly 250 ms contribute to three distinct sequences. A similar effect for Bird 3 is also observed at 550 ms and 925 ms (Figure 4G).

Across the birds the number of sequences scaled with the sample size (number of later bursts), thus from this data set there was no evident upper bound to the number of sequences. A remarkable degree of structure was observed for Bird 4, with a single 30 ms offset window accounting for 59% (10/17) of later bursts. Average numbers of bursts/offset were highest in Birds 2 and 4 (9.3 and 10 bursts/cluster, respectively), with intermediate values for Birds 1 (5.5) and 3 (5.7), and Bird 5 having the lowest numbers of bursts/offset (4). Thus, sample size was not directly related to the success of the clustering procedure. We did observe, however, that the two birds with small sample sizes had the greatest proportion of solitary bursts (on the transformed axis) (Bird 4: 0.41 (7/17), Bird 5 0.55 (7/11)). We conservatively defined sequences as having at least three bursts, but this may overly penalize the birds with the smallest sample sizes. Lowering the criterion to two bursts per sequence decreases the proportion of solitary bursts (Bird 4: 5/17; Bird 5: 3/11), bringing them more in line with the other three birds.

### Reevaluating analyses for uniform distributions

Lynch et al. (2016) described HVC PN spike bursts (combining both HVC_X_ and HVC_RA_) as forming a single continuous representation over the time course of the song motif. Clearly, our conclusions are distinct from theirs. Prior to exploring the basis for this discrepancy, we first confirmed that we could reproduce the results of Lynch et al. (2016).

Following the procedure of Lynch et al. (2016), bursts from all PNs recorded from a bird during singing were pooled and reduced to a burst time within a canonical song motif. The pooling procedure aggregates, for each bird, the burst times of all single burst PNs with all the burst times from all multiburst HVC_X_. The resulting vector of burst times is processed irrespective of the PN type, the ordinal burst position, or any other constraint. Lynch et al. (2016) graphically represented such data with the time of each burst represented on the abscissa and the ordinal position of each burst within the canonical motif represented on the ordinate (Figures 5, Birds 1– 5, bottom left insets). We term this a “burst cascade” representation to distinguish it from the neuron cascade representations of Figures 3 and 4. In a burst cascade representation, each row contains a single burst, and a column contains multiple data points only if there are two or more bursts with identical times. This depiction yields a strong tendency for typical experimental data to flow from top left to bottom right, giving the visual impression of a single, approximately straight line. The graphical decision to represent the data as spike rates tends to smear the time base of each burst causing overlap between bursts, thus reinforcing visual continuity. The degree to which burst times represented as a point process in this space are described by a continuous and uniform distribution were central questions explored in Lynch et al. (2016).

For each bird we then calculated inter-burst intervals (IBIs) from consecutive pairs of bursts as did Lynch et al. (2016), to produce a real data “population IBI”, one per bird. Histograms of these population IBI are shown for each bird in the top right insets of Figure 5. Note that the data treated in this way yield IBI histograms that all have peaks near 0 ms, tend to rapidly fall off in approximately exponential fashion, and have roughly the same attributes across the different birds. Thus, ignoring the neuronal identity of bursts in assessing the timing of information in the network destroys the actual structure present in the IBIs displayed by multiburst HVC_X_ (cf. Figure 1), that contributed 36–73% of the HVC_X_ bursts for each bird. It also obscures differences between birds. Computing similarity of all pairs of IBI histograms and all pairs of population IBI histograms (see Methods) yielded two completely non-overlapping distributions (Supplemental Table 3), with significantly different means (independent-sample t-test *p* = 2.7 x 10^-7^). Thus, the population IBI approach suppresses individual-level biological variation that exists in the data.

Following Lynch et al. (2016), we then compared the real data population IBI to 10,000 population IBIs produced from a synthetic uniform distribution over the song duration via parametric bootstrap (see Methods). These results are visualized, one per bird, as cumulative density functions (CDFs) of the real data (black lines, Figures 6A–E), the mean of the bootstrapped data (dark blue lines) and the 95% confidence intervals of the bootstrapped data (blue shading delimited by light blue lines). Our plots reasonably accurately reproduce the equivalent plots reported in Lynch et al. (2016) (see insets, Figures 6A–E) while showing additional detail since we did not apply 1 ms binning to the data as they did. Note that the real data stay confined within the confidence intervals at virtually all points for all birds. We then compared the real data CDF with the surrogate population CDFs using a Kolmogorov-Smirnov test, obtaining Bird 1 *p* = 0.640, Bird 2 *p*=0.015, Bird 3 *p*=0.421, Bird 4 *p*=0.714, Bird 5 *p*=0.811. Lynch et al. (2016) reported *p*=0.014 for Bird 2 and *p*>0.28 for the other four birds. Thus, our results closely align with those reported by Lynch et al. (2016). Furthermore, we observed that the remaining variation (for Bird 2) lies within the distribution of p-values obtained by running the overall parametric bootstrap procedure many times. Lynch et al. (2016) observed that after Bonferroni correction for five birds the K-S test is not significant for all birds, as did we. The similarity in the two sets of results gives confidence that we have reproduced the Lynch et al. (2016) method accurately. Note, however, that this method relies on negative inference – attempting to evaluate the failure to achieve statistical significance. It also assumes that the timing of spike bursts are independent from each other.

We also plotted burst cascades but with each burst time represented by a point (the weighted average of all spike times within the burst), with HVC_RA_, HVC_X_, and putative PNs distinguished by color (red, blue, and green, respectively), and with 95% confidence intervals calculated for the combined population of all PNs marked by shading around the path from first to last burst (Figure 5, Birds 1–5, central images). Considering all bursts, note that there are regions where the path of the sequential bursts exceeds the limits of the confidence intervals for each bird, in some cases substantially in magnitude (Figure 5, Bird 2) or over an extended interval of the path (Figure 5, Birds 2, 4). This contrasts with the behavior of the same data when viewed as a CDF (Figure 6), where the real data rarely or never violates the confidence intervals. This emphasizes that structure in the real data is not detected by negative inference using the K-S statistic, which relies on the CDFs and only examines the relative distributions of intervals, not their ordering.

Given these issues, we also searched for an independent biologically meaningful test to evaluate the utility of the population IBI approach. To this end, we examined the activity of RA projection neurons, that project to syringeal and respiratory brainstem motor neurons (Vicario, 1993; Wild, 1993), and are spatially organized by the syringeal muscle they innervate (Vicario, 1991). RA PNs also burst multiple times during each song motif, albeit with many more bursts than for HVC_X_. Using data from Leonardo and Fee (2005) (kindly provided by A. Leonardo) and restricting ourselves to the three Leonardo birds with the largest data sets, we conducted population IBI analysis on Leonardo Bird i9 (34 neurons, 267 bursts), numbers of bursts than those for Birds 1, 4, and 5 and comparable numbers of bursts as for Birds 2 and 3 of Lynch et al. (2016). Plotting these data in the burst cascade approach (Figure 7A, C, E) evinced clear deviations of the burst time paths from the confidence intervals, with prominent deviations for Birds i10 and i12 (Figure 7C, E). Yet, even with far greater violations of the confidence intervals for the population IBIs than seen in the HVC data, the measured CDF for each RA bird hewed closely to the corresponding CDF of the uniform distribution for that bird (Figure 7B, D, F). The K-S test verified this (Bird i9, *p*=0.129, Bird i10, *p*= 0.271, Bird i12, *p*=0.038); with Bonferroni correction for multiple comparisons all p-values are above 0.05. Yet RA activity is related to note structure (Yu and Margoliash, 1996) and variation in features of song including pitch, amplitude, and spectral entropy (Tang et al., 2014), and has been hypothesized to transform an HVC clock input into muscle coordinates (Fee et al., 2004). This suggests that the failure to reject the continuous representation null hypothesis will in general fail to yield biological insight into neural encoding mechanisms.

**Figure 7.**
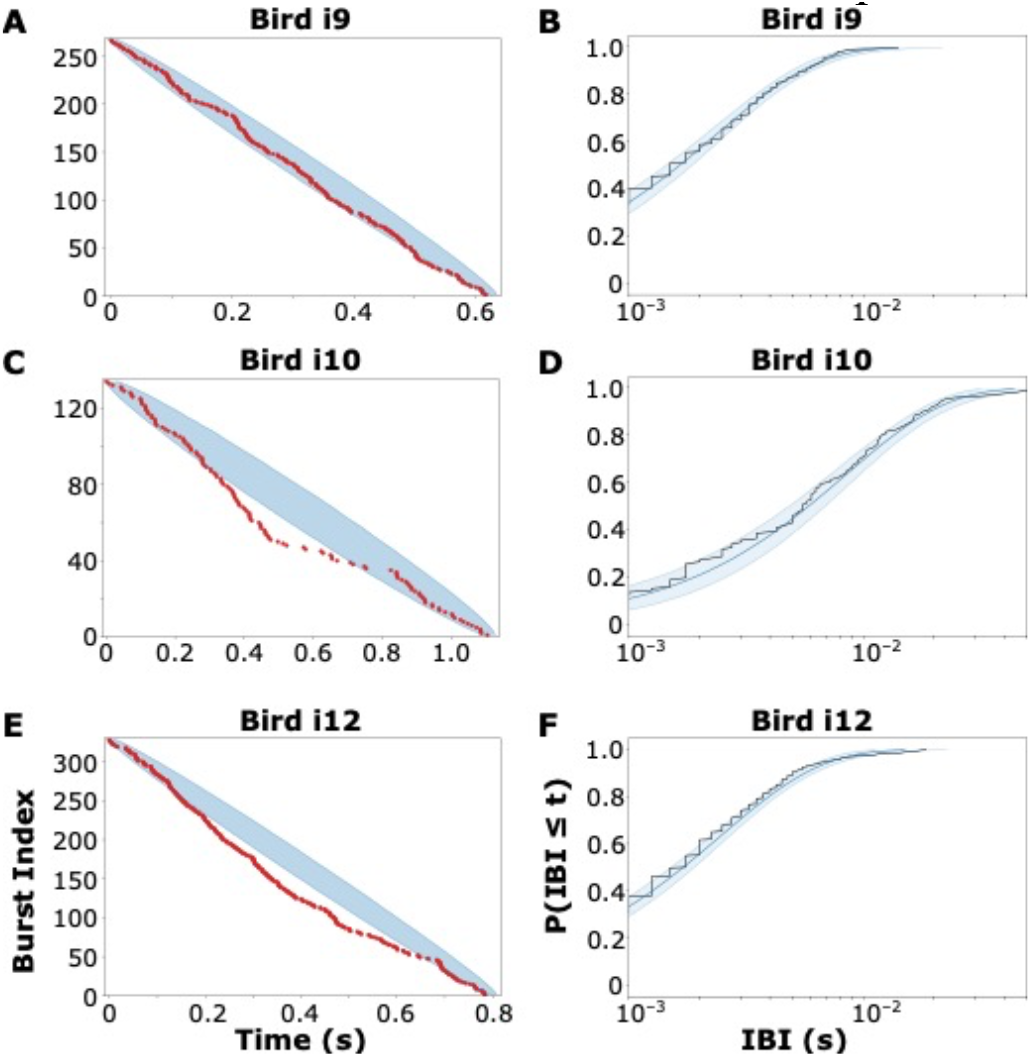
Burst cascades and IBIs of nucleus RA projection neurons recorded during singing. (A, C, E) RA PN burst times for three birds plotted as in Figure 5. (B, D, F) RA PN CDFs plotted as in Figure 6.

## Discussion

### Multiple sequences of HVC_X_ spike bursts: a model for song and error representation

Exploring the organization of HVC_X_ bursting we have identified multiple sequences of later bursts that unfold over time at different delays from the wave of first bursts. (We note that given the multiple sequences, assigning single burst neurons to the first sequence is arbitrary. Potentially, detailed functional connectivity information could resolve this issue, but this is currently unavailable.) What might the multiple sequences encode? We propose that this could be a potent organization to enhance detection of small variations in HVC activity during singing. Errors in singing could be reflected by changes in the timing and numbers of HVC_RA_ bursting at a given moment in time. The HVC_RA_ interact with HVC_X_ via local interneurons (Mooney and Prather, 2005), a disynaptic inhibitory pathway yielding song corollary discharge information in HVC_X_ (Prather et al., 2008). Within a local regime of connectivity, suppression of HVC_Int_ tonic activity could influence the timing of subsequent HVC_X_ bursting. The HVC_X_ expressing a burst timing error would burst coincidently with other HVC_X_ participating in different sequences, such as we have demonstrated here, therefore conveying the timing error laterally within the network. Such lateral interactions would involve single burst HVC_X_ as well as multiburst HVC_X_ (except those single burst HVC_X_ that occur near the beginning of song, prior to the first sequence of later bursts). Interneurons help to regulate the timing of HVC PN bursts (Amador et al., 2013; Kosche et al., 2015), and developmental learning regulates inhibition onto HVC_RA_ (Vallentin et al., 2016). Our results suggest that learning–dependent developmental changes will also be seen in the timing of HVC_X_ spike bursts.

The timing errors in the HVC_X_ potentially could also propagate forward in time (to the next bursts of those neurons). This depends on a multitude of factors, both of network activity and HVC_X_ cellular mechanisms. The latter could include IPs that influence burst timing (see below).

Whatever the sources of timing errors, and whether they arise from direct effects on HVC_RA_ or from effects in RA, the brainstem, or the basal ganglia pathway (Hamaguchi et al., 2014), error in motor output would be conveyed back to HVC through multiple pathways including via auditory feedback (Konishi, 1965). In an experiment where zebra finch were presented with a continuous delayed auditory feedback (cDAF), song was rapidly and potently disrupted (Fukushima and Margoliash, 2015). Presentation of cDAF also disrupted HVC_X_ IP homogeneity, in a dose-dependent fashion. Changes in the IPs of HVC_X_ were detected in as little as 4 hours after the onset of cDAF, the shortest interval attempted (Daou and Margoliash, 2020). These results indicate that auditory feedback error (or motor error from auditory feedback error) can rapidly drive changes in HVC_X_ IP homogeneity, representing an error signal. Examining the changes in HVC that accompany cDAF singing can help to elaborate this model.

Biological systems can experience substantial delay in feedback relative the motor event giving rise to that feedback. Typically, this problem is posed it terms of how feedback can “find” and alter the neurons whose motor-related activity gave rise to that feedback earlier in time. Instead, we envision the problem as mapping a sequence of feedback events onto a sequence of delayed copies of motor-related events (multiple spike bursts within individual neurons). The organization of HVC_X_ spike bursts into multiple sequences preserves the temporal precision and sparseness of individual HVC_X_ bursts while organizing them into a dense population representation at each moment in time. This structure could also accommodate feedback signals from respiratory, auditory and somatosensory activity representing information about singing behavior that are transmitted to HVC (Wild, 1994; Suthers et al., 2002; Ashmore et al., 2005; Akutagawa and Konishi, 2010) presumably with different delays. A given feedback signal would synapse into a local circuit (including HVC_X_ and HVC interneurons) at the appropriate moment in time that compensates for the delay associated with that input, allowing it to interact with corollary discharge information in the later sequences. Perhaps during development feedback signals project widely through HVC, allowing local networks within HVC to preferentially enhance connections based on temporal coincidence (pruning non-coincident connections), resulting in tuning to specific delays. Thus, our results suggest that sparse, structured HVC_X_ bursting effectively carries an efference copy signal, representing a biologically plausible solution to the so-called temporal credit assignment problem.

Finally, the projection of HVC_X_ would transmit to the basal ganglia pathway a normal signal (no error) or an error signal, with the signal converted into motor commands instructing RA activity (Giret et al., 2014). The signal might be evaluated in the basal ganglia by assessing the coincidence of populations of HVC_X_ bursts occurring over small intervals of time.

### Spike rebound excitation as a candidate mechanism for network rhythmic bursting

We propose that the timing of HVC_X_ spike bursts depends both on the pattern of connections in HVC and on HVC_X_ IPs through the mechanism of spike rebound excitation. In spike rebound excitation, the timing of spike bursts are regulated by IPs that influence the duration and magnitude of hyperpolarizing membrane excursions interacting with release from inhibition (a consequence of changes in network activity) that rapidly triggers a spike burst.

In vitro recordings demonstrate that HVC_X_ express spike rebound excitation (Daou et al., 2013). In HVC_X_ this is mediated by the expected complement of ion channels, including hyperpolarization-activated cyclic nucleotide-gated channels (*I_h_*), T-type calcium channels (*I_ca-T_*), and small conductance calcium-activated potassium channel (*I_SK_*) (Daou et al., 2013). Previous results indicate that relative minima of HVC_Int_ occur in temporal relation to spike bursts of HVC_X_ (Amador et al., 2013) and HVC_RA_ (Kosche et al., 2015) during singing. These are the most fundamental set of features required to support the spike rebound excitation mechanism, but absent additional constraints, demonstrating the action of these features in behaving animals can be difficult (e.g., making whole cell patch recordings in singing birds).

The organization of the HVC_X_ in zebra finches provides for such additional constraints (Daou and Margoliash, 2020). Recent results with in vitro whole cell patch recordings in brain slices from adult birds demonstrate that all the HVC_X_ in a given animal tend to have similar spike waveform shapes and timing of spikes in response to canonical depolarizing and hyperpolarizing current injections. Following Hodgkin Huxley modeling, this yields similar estimates for each of five ion current magnitudes (I_Na_, I_K_, I_Ca-T_, Ih, and I_SK_) in different neurons within the same animal. Different individuals have different combinations of IPs (except birds singing similar songs such as sibling birds). Differences in the magnitudes of I_Ca-T_, I_h_, and I_SK_ could induce substantial effects on the timing of spike rebounds bursts. Correspondingly, systematic differences among individual birds in these IPs could induce systematic differences in the timing of spike rebound bursts. This argues a for a connection between the IP homogeneity seen in vitro and the multiple spike burst sequences observed in vivo. We envision that this pattern of organization will vary among songbird species depending on the regularity of singing structure.

Given that the homogeneity of HVC_X_ IPs in adults is sensitive to experimentally-induced errors in singing (Daou and Margoliash, 2020), that HVC_X_ IPs are developmentally regulated (Ross et al., 2017; Ross et al., 2019; Daou and Margoliash, 2020), and that juvenile zebra finches in the plastic stage of song development show differences in their IPs and far less uniformity in IPs as compared with adults (Ross et al., 2017; Daou and Margoliash, 2020), this adds confidence that HVC_X_ IPs are closely tied to the song of the bird. Nonetheless, we have not yet associated specific IP values with specific features of song or specific patterns of spike burst timing during singing. These represent attractive future directions.

### Reevaluating the continuous representations hypothesis for HVC

The clock model envisions a single continuous representation of burst times, with recent papers combining both HVC_X_ and HVC_RA_ into a single sequence (e.g., Lynch et al., 2016; Picardo et al., 2016). Feedforward excitatory connections are envisioned as the mechanism to sustain propagation of bursts down the chain. The activity of each HVC PN is said to explicitly represent time itself, hence these are “clock” models. The bursts of multiburst HVC_X_ represent a challenge, however, since under the feedforward concept the functional connections each HVC PN makes will presumably be invariant to which sequence it is participating in, yet the sequences represent different moments in time. In contrast, our model proposes a simple mechanism for implementing dynamics in functional network connectivity.

As well as conceptual issues there are statistical ones. Lynch et al. (2016) treated HVC PN burst times as independent (Figure 5, Birds 1–5, bottom insets). A similar assumption was made by Picardo et al. (2016) which used optogenetic techniques to image large numbers of HVC PNs neurons during singing. This included recording from multiburst neurons but in that paper too the timing of each burst was treated as an independent event (and multiple bursts were not considered in the analysis). Our results demonstrating rhythmicity of HVC_X_ bursting renders the assumption of spike burst independence untenable and invalidates the basic statistical treatment of data in those papers. Both Lynch et al. (2016) and Picardo et al. (2016) concluded that HVC PN are organized in a single continuous representation across the song motif. Our results refute this for HVC_X_ PN and emphasize that the neuronal identity of bursts has biological as well as statistical significance.

Central to the statistical testing proposed for evaluating a continuous representation is attempting to support a model by appealing to the failure to reject the null hypothesis. Innumerable forms of structured data may hide behind such a failure, including the results we reported. Furthermore, we showed that the KS statistic as applied to populations of IBIs is weak and uninformative, failing to detect evident non-uniform distributions in the sequence of IBIs. Yet the clock model envisions for each bird a specific sequence of IBIs. The population IBI concept orders bursts independent of the neurons or ordinal positions of the bursts. Whether this has utility remains unknown. Instead, we found structure by respecting the neuronal identity and sequential order of the bursts.

Finally, here and in other recent papers very small samples of HVC_RA_ have been “hitched” onto much larger samples of HVC_X_ (Okubo et al., 2015; Lynch et al., 2016) for analysis, or small numbers of intracellularly recorded HVC_RA_ were analyzed (Picardo et al., 2016). We fully acknowledge all these remarkable recordings during singing are exceptionally difficult to achieve, and represent a tour de force. Nonetheless, when small samples of HVC_RA_ are considered in isolation, they define an almost limitless space of potential paths through the motif (Figure 5, bottom right panel). Thus, they offer effectively no information about the sequential organization of populations of HVC_RA_ in relation to singing. In contrast, the optical recordings of Picardo et al. (2016) included presumptive large samples of both HVC_RA_ and HVC_X_, but for reasons of technical limitations they could not distinguish between the two classes of PN, and therefore also provided no resolution to this problem. Thus, we have shown that the temporal organization of HVC_X_ bursts are not consistent with the continuous representation clock model, and it continues to remain entirely in the realm of speculation whether HVC_RA_ are.

## Methods

### Data collection

These data were collected by the authors of Lynch et al. (2016) and provided by them at our request. We restate the salient details here; for more detail on collection, refer to their methods. All HVC projection neuron data were collected using microdrive-mounted high-impedance single electrodes surgically implanted into HVC of male zebra finches. Electrophysiological recordings were made while the birds sang in the presence of a female zebra finch (directed song) and neurons were identified as interneurons or PNs according to firing properties. PNs were classified as HVC_RA_ or HVC_X_ based on antidromic stimulation using electrodes implanted into nucleus RA or Area X, or were classified as “putative PNs” if they demonstrated song-related firing patterns characteristic of PNs but could not be definitively identified by antidromic stimulation.

All RA projection neuron data were collected similarly, with projection neurons identified by their firing properties. These data were previously reported in Leonardo & Fee (2005) and were provided to us by Anthony Leonardo. For more detail, refer to that paper’s methods.

### Data received

We received both spike times and inferred burst times from the Lynch et al. (2016) authors. In both cases, the values provided were those obtained after having been warped to a canonical motif for each bird (piecewise linearly by syllable start and stop times, following Glaze and Troyer 2006).

Lynch et al. (2016) determined burst times by producing 1 ms-binned spike rates pooled over all motif renditions, smoothing by convolution with a 9ms square window, and identifying candidate burst onset and offset times using a 10Hz threshold. Only candidate windows which contained spikes on greater than 50% of all song renditions were considered. Burst times were calculated from these windows by taking the mean spike time within the window.

In all analyses involving burst timing, we used the burst times provided to us, rather than recalculating them ourselves (beyond confirming that we were able to), to maintain continuity with the original data and analyses.

### Identification of PNs

We treated all multiburst neurons as HVC_X_. Thus, we include the three multiburst HVC_RA_ from the Lynch et al. (2016) data (one neuron in each of Birds 2, 4, and 5) given that such HVC PNs are not described elsewhere and we cannot exclude the possibility that like canonical HVC_X_ these neurons also project to Area X (the basal ganglia) (see Benezra et al. (2018)). Any bias towards one or the other class of identified PN in the putative PNs is not known.

### Application of Lynch analysis to RA data

The RA data provided by A. Leonardo included spike rates in 0.5 ms bins, averaged over all motif renditions (after piecewise linear warping to a canonical motif). As the firing properties of RA PNs differ from those of HVC PNs, we could not apply the burst identification method used by Lynch et al. (2016). Following Leonardo & Fee (2005) we identified burst onsets and offsets with a 125 Hz threshold on this spike rate.

Surrogate random data were generated largely following the procedure described above, except that the prohibition on bursts from a single neuron closer together than 9 ms was removed. Otherwise, all analyses performed on these data were identical to those for the HVC PN data.

### Surrogate uniform distributions

Surrogate uniform distributions for comparison with the real data were generated by parametric bootstrap as follows: to generate a single surrogate dataset for one bird, new burst times were generated for each neuron from that bird, matching the number of bursts that neuron produced. These burst times were drawn uniformly from an interval with length equal to that of the bird’s canonical motif, with the requirement that for a given neuron, no two bursts occur within 9ms of one another (as the burst identification analysis described above precludes this possibility in the real data). Burst times were generated independently across neurons. Population IBIs were generated as for the real data. (Lynch et al. (2016), personal communication with Galen Lynch).

For each bird, 10,000 of these surrogate datasets were generated.

### Population IBI CDFs and Kolmogorov-Smirnov tests

The primary statistical test performed on the population IBIs was a two-sample Kolmogorov-Smirnov (K-S) test between the real data and the median of the 10,000 surrogate uniform distributions for each bird. The K-S statistic generated was compared with the distribution of K-S statistics calculated between the median and each of the 10,000 surrogate distributions. The p-values quoted in the results express the proportion of this bootstrapped distribution which is greater than or equal to the K-S statistic obtained between the real data and the median of the surrogates (after a small-sample correction). The K-S test was applied using the ks_2samp method from the Python scipy.stats library (here and elsewhere in our analyses).

### Multiburst neuron burst timing analysis

The multiburst neuron IBI distributions were generated by first calculating the IBIs between the bursts for each neuron, and then pooled into a single distribution per bird. That is, if a bird has two multiburst neurons, each of which fires two bursts, then each neuron will contribute one IBI to that bird’s multiburst neuron IBI distribution. For all histograms displaying these distributions a bin size of 10 ms was used.

To quantify histogram similarity, for each bird we constructed 10-bin histograms with the ordinate normalized by the peak height of each histogram, both for standard IBI and population IBI representations. We compared pairs of histograms by computing an RMS value summed over the difference between bin 1 pairs, bin 2 pairs, etc. The smaller the summed number, the more similar on average are the two histograms being compared.

### Set cover analysis

The one-dimensional geometric set cover problem can be solved greedily by sorting the sets by their central point and sweeping across the entire interval in one direction, and for every uncaptured burst encountered, collecting the set that contains that burst on its trailing edge and marking all bursts within that set as captured. This solution will be optimal with respect to the total number of sets (or offset times) required to capture all of a bird’s later bursts. (This algorithm is usually discussed without attribution, given its simplicity, e.g. Agarwal et al (2020).)

The alternative algorithm we used to maximize the area under the capture curve is also simple: collect sets (and mark the bursts they contain as captured) in decreasing order of the number of uncaptured bursts contained in the set. This algorithm will produce solutions with more sets that capture a large number of bursts, at the cost of a longer “tail” of sets which only capture one burst.

## Code availability, quantification

All code and statistical analyses will be made available upon publication on GitHub (exact location TBD). We used Python, various Python libraries, and Jupyter notebooks. The complete list and version numbers are found in the GitHub repository.

## Acknowledgements

We are grateful to Gabriel B. Mindlin, Jason N. MacLean, Sliman J. Bensmaia, Nicholas G. Hatsopoulos, Stephanie E. Palmer, and Arij Daou for comments on earlier versions of the manuscript. Supported by NSF NCS grant 1835181 and NINDS grant UF1NS115821-01.

---

**Supplemental Figure 1.**
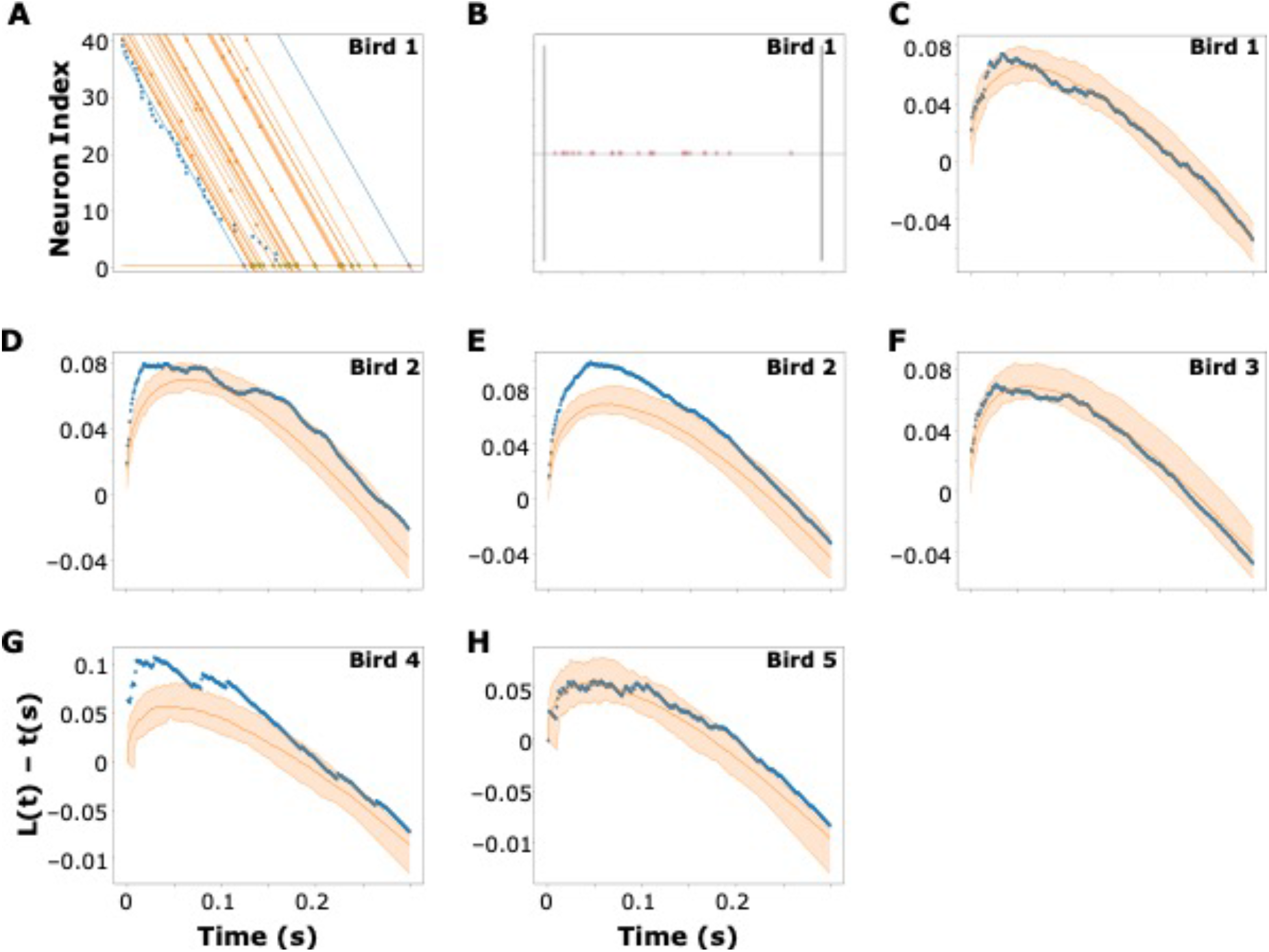
Linear Fits of Later Bursts in Neuron Cascade Representations Demonstrate Clustering (A) Starting with the neuron cascade representation for Bird 1 (Figure 3A), a line (left blue line) was defined as the best fit to all the initial bursts. Parallel lines (in orange) were then drawn through each of the subsequent bursts except the line for the last burst is in blue. An axis normal to the lines was then defined. While the normal axis appears graphically to be far from normal this is due to the different axis scales (roughly 100 to 1). The intersection points of the blue lines with the normal line are depicted as red circles and define the bounds of possible projections of later bursts onto the normal line. (B) The normal axis line from (A) with the burst intersection points and bounds (vertical lines). Note many bursts appear to cluster together. (C—G) Statistical analysis using Ripley’s L statistic of clustering for each piecewise fit of initial bursts for each of the five birds. (Only Bird 2 had two piecewise fits associated with significant numbers of subsequent bursts and so contributes two plots.) The observed clustering along the normal axis was compared with clustering in synthetic data that were produced so as to maintain the locations of each neuron’s first bursts, but to uniformly randomly place its subsequent bursts between its first burst and the end of the canonical song motif (maintaining a 9 ms interval between any two bursts, per Lynch et al. (2016)). Short-range clustering on this axis indicates that the later bursts of multiburst neurons tend to describe sequences during singing that proceed at the same rate as the sequence described by the first bursts of these neurons. For Birds 1–4 the normal-axis intersection clustering exceeds the 95% confidence interval calculated via the bootstrap procedure at short cluster distances. This was limited to brief intervals for Birds 1 and 3 (C and F, respectively), but with substantial departures from the 95% confidence intervals for Bird 2 (both fits) (D and E) and Bird 4 (G). Bursts from Bird 5 (H) exceed the confidence interval only for clustering at long cluster distances; however, given that Bird 5 had very few later bursts (n = 11 later bursts; Supplemental Table 1) it exhibited wide confidence intervals at short cluster distances. Bird 4, with the next-fewest later bursts (n = 17; Supplemental Table 1), was the only other bird to share these features of the shape of the confidence intervals: wide intervals at short cluster distance and exceeding the confidence interval at long cluster distance.

**Supplemental Table 1.**
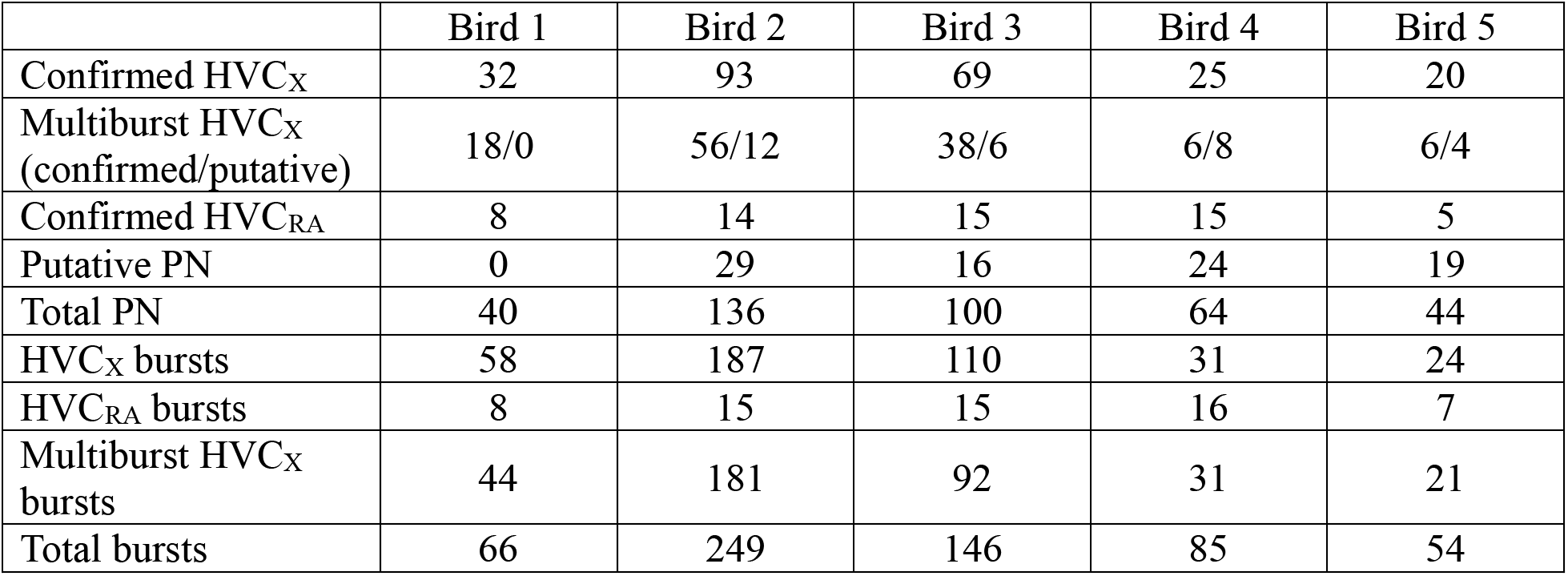
Numbers of different classes and spike bursts of HVC projection neurons.

|  | Bird 1 | Bird 2 | Bird 3 | Bird 4 | Bird 5 |
| --- | --- | --- | --- | --- | --- |
| Confirmed HVC<sub>X</sub> | 32 | 93 | 69 | 25 | 20 |
| Multiburst HVC<sub>X</sub><br>(confirmed/putative) | 18/0 | 56/12 | 38/6 | 6/8 | 6/4 |
| Confirmed HVC<sub>RA</sub> | 8 | 14 | 15 | 15 | 5 |
| Putative PN | 0 | 29 | 16 | 24 | 19 |
| Total PN | 40 | 136 | 100 | 64 | 44 |
| HVC<sub>X</sub> bursts | 58 | 187 | 110 | 31 | 24 |
| HVC<sub>RA</sub> bursts | 8 | 15 | 15 | 16 | 7 |
| Multiburst HVC<sub>X</sub><br>bursts | 44 | 181 | 92 | 31 | 21 |
| Total bursts | 66 | 249 | 146 | 85 | 54 |

**Supplemental Table 2.** Testing for bird-specific structure in the autocorrelograms. K-S test p-values for comparisons of distributions of neuron-by-neuron autocorrelogram dot products between all pairs of birds. The matrix is not symmetric because the distribution of dot products between a pair of birds’ neurons can be compared with each bird’s distribution of its own neurons dotted with one another.

| Bird | 1 | 2 | 3 | 4 | 5 |
| --- | --- | --- | --- | --- | --- |
| 1 |  | 1.31e-36 | 2.22e-16 | 2.03e-29 | 1.12e-14 |
| 2 | 0.00135 |  | 1.81e-45 | 1.83e-25 | 2.47e-08 |
| 3 | 2.08e-09 | 1.9e-28 |  | 1.21e-14 | 0.0147 |
| 4 | 2.66e-15 | 6.66e-16 | 2.61e-10 |  | 0.523 |
| 5 | 1.28e-14 | 6.53e-29 | 0.00167 | 0.000192 |  |

**Supplemental Table 3.**
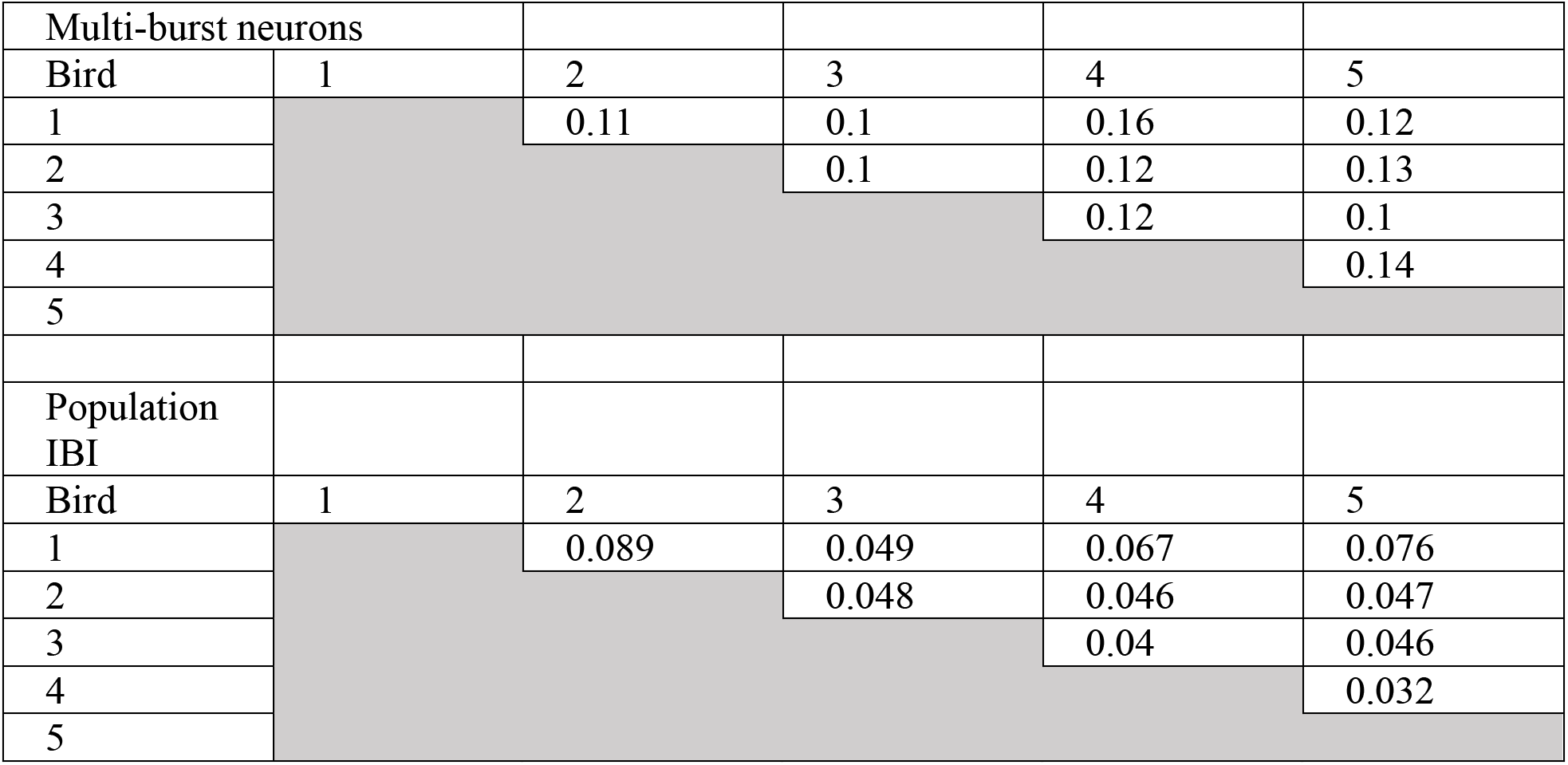
Comparing structure of multiburst IBI histograms and population IBI histograms. Root mean square of bin differences from multiburst neuron IBI histograms (top) and population IBI histograms (bottom).

